# Range-wide responses to an extreme heat event in *Mimulus cardinalis*

**DOI:** 10.1101/2025.05.01.651703

**Authors:** Lucas J. Albano, Robin A. Bingham, Sulma Correa, Catherine G. Laufenberg, Cristina Payst, Christopher D. Muir, Seema Nayan Sheth

**Author notes:** **Correspondence Author**: Lucas J. Albano, Current address: Department of Plant & Microbial Biology, North Carolina State University, 100 Derieux Place, Raleigh, NC, USA, 27605.

## Abstract

**Premise:** Extreme events are an understudied aspect of ongoing anthropogenic climate change that could play a disproportionate role in the threat that rapid environmental shifts pose to natural populations.

**Methods:** We used a short-term heatwave treatment in controlled growth chamber environments to study evolutionary responses in *Mimulus cardinalis* populations to a recent repeated extreme heat event. Individuals from before (ancestors) and after (descendants) this extreme heat event were resurrected from seeds harvested from six populations across the range of *M. cardinalis*, and their physiological, performance, and functional responses (stomatal conductance, leaf temperature difference, photosystem II efficiency, relative growth rate, specific leaf area, and leaf dry matter content) to the heatwave treatment were assessed.

**Key Results:** Our results allow for two main conclusions: 1) plants responded favorably to the heatwave treatment in terms of their overall performance and the magnitude of response was generally greatest among trailing-edge populations, and 2) despite some evidence of divergent evolutionary responses among regions to the recent natural extreme heat event, we did not find adaptive evolution that affected how *M. cardinalis* populations responded to the heatwave treatment.

**Conclusions:** These results demonstrate that many *M. cardinalis* populations may be residing in natural environments that are below their optimum average temperature, indicating that *M. cardinalis* populations could exhibit resiliency to future warming. However, limited evolutionary responses in *M. cardinalis* to the recent extreme heatwave could still indicate that there is potential for future vulnerability to extreme climate events of increased intensity, frequency, and duration in the future.

## Introduction

Ongoing climate change is associated with an increase in the frequency, duration, and intensity of extreme events, such as floods, storms, droughts, and heatwaves (IPCC, 2022). Extreme climate events (i.e., those with a ≤ 5% probability of occurring; Smith, 2011) are recognized as critical components of climate change and can be important selective forces for natural populations (Gutschick and BassiriRad, 2003; Grant et al., 2017). For example, temperature extremes are expected to have severe and widespread ecological impacts (Smith, 2011; Ruthrof et al., 2018; IPCC, 2022), yet a disproportionate number of previous studies have focused solely on impacts of the mean temperature increases associated with climate change (Breshears et al., 2021). In terrestrial systems, such temperature extremes can be defined as “heatwaves” when three or more consecutive days experience maximum temperatures above the 90th percentile for a particular location at a particular time (Perkins and Alexander, 2013). Global climate models project not only an increase in absolute maximum temperatures, but also increases in the intensity, frequency, duration, and geographic extent of heatwaves (Meehl and Tebaldi, 2004; Seneviratne et al., 2012; Coumou and Robinson, 2013; Coumou et al., 2013; Perkins-Kirkpatrick and Gibson, 2017; Guerreiro et al., 2018; IPCC, 2022). Heatwaves in North America have already become more extreme between 1961 and 2021, with increases in average frequency (from two per year to six per year), duration (from 3 days to 4 days), and intensity (from 2.0°C to 2.3°C) during that period (NOAA, 2024). Therefore, both past data and future projections point to an urgent need to improve our understanding of population vulnerability to heatwaves and the potential for adaptive evolution in response to extreme climate events.

In general, populations exhibit a breadth of thermal performance (i.e., a range of temperatures at which they are considered to be reasonably successful) as well as a thermal optimum (i.e., a specific temperature at which they are most successful; Huey and Stevenson, 1979; Huey and Kingsolver, 1989). Populations exposed to temperatures outside of their thermal performance breadth could experience a reduction in performance and fitness (Huey and Stevenson, 1979; Huey and Kingsolver, 1989). In plants, the thermal safety margin describes the difference between the temperature at which leaves begin to lose functionality (i.e. T_50_, or the temperature at which photosystem II activity falls below 50%) and the actual leaf temperature (Ahrens et al., 2021; Cook et al., 2021; Kitudom et al., 2022). High temperatures (i.e., those exceeding the thermal performance breadth for a particular plant population, which is more likely to occur under narrower thermal safety margins) can result in detrimental effects on numerous physiological processes, including stomatal closure, inhibition of photosynthetic machinery, accumulation of reactive oxygen species, and/or slowing of nitrogen metabolism (Hasanuzzaman et al., 2013; Feller and Vaseva, 2014; Teskey et al., 2015; dos Santos et al., 2022). Such extreme heat conditions could negatively impact plant survival and reproduction to the point that populations may be unable to persist in the face of ongoing climate change (Gutschick and BassiriRad, 2003; Teskey et al., 2015). Extreme heat also serves as a strong selection pressure on plant populations, with individuals that are more successful in avoiding or tolerating the detrimental effects of extreme heat being more likely to survive and reproduce (Gutschick and BassiriRad, 2003; Teskey et al., 2015; Grant et al., 2017). Genetic variation in mechanisms of resistance to extreme heat within a plant population presents the possibility of adaptive evolution as an overarching means of population persistence in the face of increasingly detrimental effects of extreme heat events (Orsenigo et al., 2014; Lancaster and Humphreys, 2020; Martin et al., 2023).

While adaptive evolution can be a key mechanism by which populations cope with climate extremes (Grant et al., 2017), we lack a clear understanding of whether populations across a species’ range can adapt fast enough to persist in the face of more frequent and intense heatwaves. Populations across species’ ranges may vary in their ability to adapt to extreme climate events based on range limit theory (Peischl et al., 2015; Sexton et al., 2016; Lancaster and Humphreys, 2020). While trailing-edge populations could better tolerate extreme heat, due to greater historical exposure to higher temperatures (Angert et al., 2011; King et al., 2019; Chiono and Paul, 2023), climate warming could reduce trailing-edge population sizes and adaptive potential (Jump et al., 2006, 2010; but see Sheth and Angert, 2016). Conversely, under equilibrial range limit theory, populations located at the range center are expected to harbor the greatest genetic variation and adaptive potential (Jump and Peñuelas, 2005; Guo, 2012; Kremer et al., 2012; Latron et al., 2020). An influx of range-center migrants at the leading edge as the climate warms could therefore boost genetic variation and adaptive potential at the leading edge (Jump and Peñuelas, 2005; Guo, 2012; Kremer et al., 2012; Latron et al., 2020). However, conditions of range limit theory are generally not in equilibrium due to shifting environments and populations, and recent theory suggests more nuanced trends of genetic variation and adaptive potential across a species’ range (Polechová, in review). For example, range-edge populations may be locally adapted to more variable climate conditions and therefore could be integral to a species’ ability to withstand the effects of climate change across its range (Stevens, 1989; Rehm et al., 2015; King et al., 2019; Preston et al., 2022). Comparing responses to a historic heatwave in populations across a species’ range can therefore provide geographical context to our understanding of how natural populations respond to extreme climate events and their ecological consequences.

A resurrection study is a powerful approach to assess evolutionary responses in natural populations, by simultaneously growing and comparing seeds harvested from different timepoints (i.e., ancestors versus descendants; Franks et al., 2007, 2008, 2018). We implemented a resurrection growth chamber experiment comparing responses to a heatwave treatment between ancestral (2010) and descendant (2017) cohorts from populations across the geographic range of the scarlet monkeyflower, *Mimulus cardinalis* (syn. *Erythranthe cardinalis*; Lowry et al., 2019; Sheth et al., in review). Specifically, we used seeds collected from six *M. cardinalis* populations (two leading-edge, two range-center, and two trailing-edge) both before (ancestors) and after (descendants) a multi-year period of extreme heat and drought in western North America (Diffenbaugh et al., 2015; Robeson, 2015). Populations across the range of *M. cardinalis* may vary in their ability to respond to heatwaves based on not only the magnitude of temperature extremes they have experienced, but also their possession of mechanisms to withstand extreme heat (Angert et al., 2011; King et al., 2019; Lancaster and Humphreys, 2020; Kitudom et al., 2022; Chiono and Paul, 2023). Previous work found rapid responses to artificial selection on flowering phenology in trailing-edge populations, as well as correlated changes in leaf functional traits, which is contrary to expectations based on range limit theory (Sheth and Angert, 2016). In this study, we are interested in testing the effects of heatwaves on physiological, performance, and functional traits. For example, stomatal conductance of water (*g*_sw_) is an important measure of the rate of gas exchange in plant tissue, with high *g*_sw_ suggesting the potential for high rates of photosynthesis and transpiration, and low *g*_sw_ suggesting plants may be conserving water and photosynthesizing more slowly (Turner, 1991; Urban et al., 2017). *g*_sw_ is related to leaf temperature difference (air temperature − leaf temperature), which indicates not only a physical cooling effect based on heat loss through transpiration, but also a potential adaptive response to maintain leaves at a temperature that is more ideal for photosynthesis (Farquhar and Sharkey, 1982; Teskey et al., 2015; Griffani et al., 2024). Similarly, high photosystem II efficiency (Φ_PSII_, or the proportion of light absorbed by photosystem II that is used for photochemistry) indicates a healthy plant that is conducting photosynthesis efficiently and has the potential for a high relative growth rate, while low Φ_PSII_ indicates potential plant stress and likely lower relative growth rate (Bilger et al., 1995; Neri et al., 2024). Specific leaf area (SLA) and leaf dry matter content (LDMC) are widely used metrics of plant response to environmental conditions (Pérez-Harguindeguy et al., 2013). In general, plants in hotter, drier environments produce leaves with lower SLA and higher LDMC, and have been reported to better recover photosynthetic activity after heat stress (Knight and Ackerly, 2003; Reich, 2014; Leigh et al., 2017; Liu et al., 2023). Using these traits to compare responses of resurrected populations to a heatwave treatment can therefore elucidate potential patterns of evolution to extreme climate across the species range.

Our objectives were to 1) quantify responses in plant physiological, performance, and functional traits to a heatwave treatment and determine if those responses varied among populations from across the range of *M. cardinalis*, and 2) characterize differences in response to a heatwave treatment between 2010 ancestors and 2017 descendants, indicative of an evolutionary response to the recent heat and drought event experienced by natural *M. cardinalis* populations. To address these objectives, we quantified numerous traits expected to be involved in plant response to the surrounding environment, namely *g*_sw_, leaf temperature difference, Φ_PSII_, relative growth rate in leaf number (RGR), SLA, and LDMC (Pérez-Harguindeguy et al., 2013; Reich, 2014). We predicted reduced plant performance under the heatwave treatment and that the magnitude of these responses would be greatest among the least warm-adapted (leading-edge) populations, based on their lower historical exposure to high temperatures. If adaptive evolution has occurred due to the recent heat and drought event, we predicted that descendants would exhibit shifts in plant physiological, performance, and functional traits compared to ancestors.

However, under a scenario of adaptive evolution in response to extreme heat, for descendants relative to ancestors, we expected to observe shallower responses between the heatwave treatment and the control, indicating greater resistance to the extreme heat. Furthermore, since leading-edge and range-center populations experienced the greatest magnitude of warming between 2010 and 2017, we expected to observe the smallest reduction in performance (descendants relative to ancestors) to occur in these populations compared to trailing-edge populations. Our results contribute to our understanding of how plant populations adapt to changing environments over space and time, and whether evolution can rescue populations that are declining due to climate change and the associated increases in extreme climate events.

## Methods

### Study system and source populations

*Mimulus cardinalis* is a perennial herb ranging from southern Oregon, USA (leading range edge) to northern Baja California, Mexico (trailing range edge; **Figure 1**). It occurs in seasonally wet riparian zones below 2400 m throughout this region. The subject of numerous previous studies, *M. cardinalis* is a model system for the study of evolutionary, physiological, and demographic responses to climate change (Angert and Schemske, 2005; Angert et al., 2011; Sheth and Angert, 2016, 2018; Muir and Angert, 2017; Anstett et al., 2021; Branch et al., 2024; Anstett et al., in review; Sheth et al., in review). All plant material used for this study originated from a collection of *M. cardinalis* seeds that were previously harvested from each of two leading-edge (N1 and N2), two range-center (C1 and C2), and two trailing-edge (S1 and S2) populations, as described in Sheth and Angert (2016). From 2010 to 2014, population growth rates for 19 of 32 *M. cardinalis* populations decreased (including N1, C2, and S1), with the lowest probability of survival occurring at both the leading and trailing range edges (Sheth and Angert, 2018). Between 2010 and 2017, the leading-edge (N1 and N2) and range-center (C1 and C2) source populations experienced the greatest increase in temperature compared to previous years, while temperatures at the trailing edge became less variable (**Figure 1**). For each population, seeds were collected from 80-100 individuals (representing unique maternal families) in the fall of 2010 (ancestral cohort) and in the fall of 2017 (descendant cohort). A refresher generation was created by crossing plants within each population and cohort, generating 18 full-sib seed families per population and cohort combination in order to reduce maternal and seed age effects, as described in Wooliver et al. (2020).

**Figure 1.**
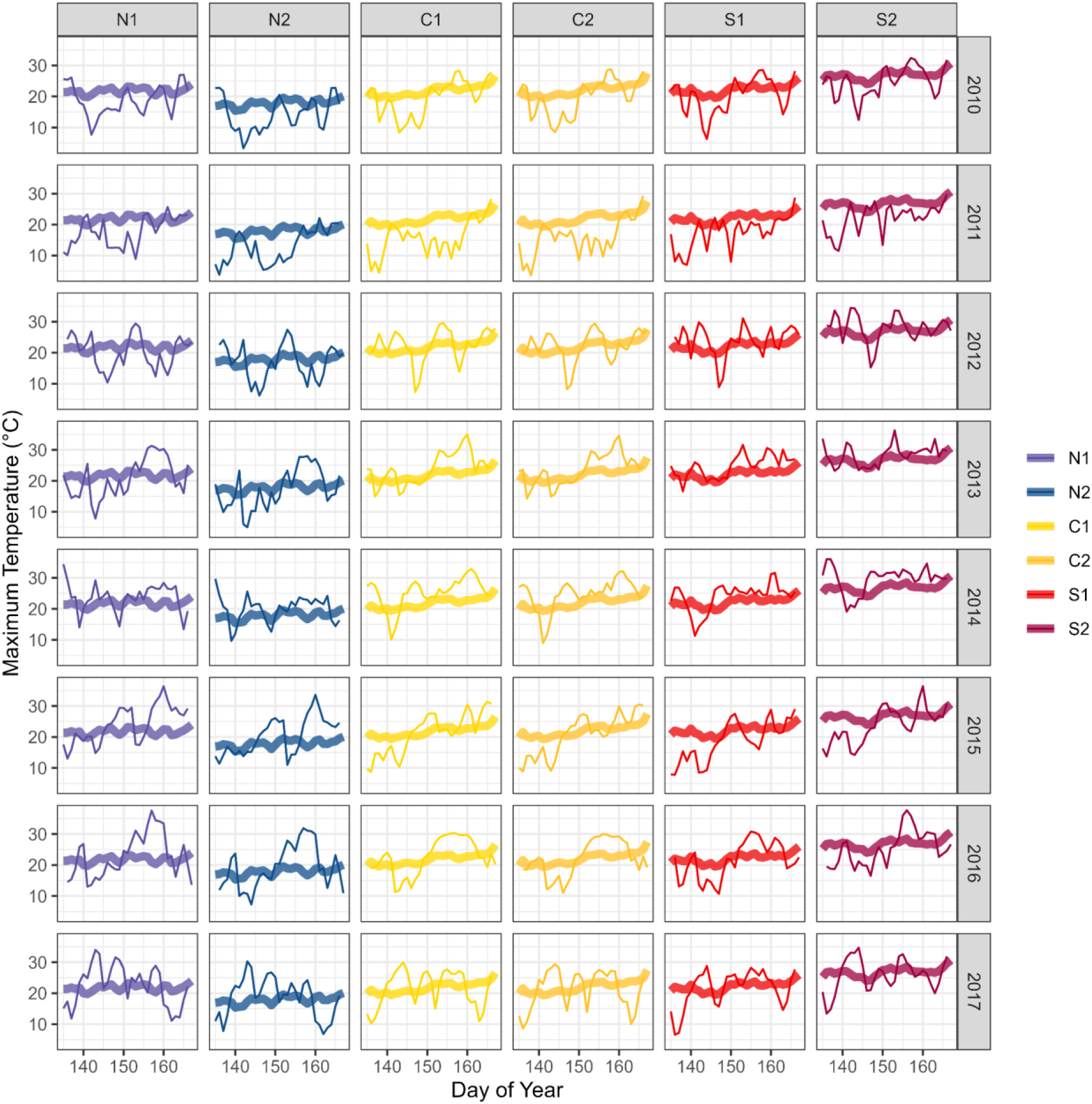
Historical and recent heatwave temperatures for the six focal *M. cardinalis* populations used in our study. The bold lines for each of the two leading-edge (N1 and N2), two range-center (C1 and C2), and two trailing-edge (S1 and S2) source *M. cardinalis* populations represent historical (1990-2010) daily average maximum temperatures from May 15 to June 15 each year. Thin lines represent daily maximum temperatures from May 15 to June 15 of each year from 2010 (ancestral cohort) through 2017 (descendant cohort), including all individual years of the recent repeated extreme heat event. The most extreme temperature across all years and populations (population S2; June 4, 2016/day 155; 37.7°C) guided the temperature regime for the heatwave treatment in our study. All climate data was extracted from PRISM (PRISM Climate Data; (PRISM Group, 2014).

### Experimental design

To determine whether *M. cardinalis* has evolved in response to recent extreme events as a result of climate change, we conducted a resurrection experiment in growth chambers using the refresher generations of ancestral and descendant seed families from the six selected source populations. We planted seeds into twenty-four 72-cell plug trays (one seed per cell) filled with autoclaved Sunshine Mix #4 Aggregate Plus (Sun Gro Horticulture, Agawam, MA, USA), topped with a thin layer of autoclaved cement sand (Southern Products & Silica Co., Hoffman, NC, USA) and thoroughly misted with water. Planting was staggered by tray and occurred from February 21 through March 6, 2024. To reduce the effect of competitive interactions arising from among-region variation in germination time and growth rate, seeds from each region were planted in separate trays. Each tray contained one plant from each of the 18 seed families per population and cohort combination (18 seed families × 2 populations per region × 2 cohorts per population = 72 plants per tray, arranged randomly). Seedlings were established in a common walk-in growth chamber environment under fluorescent lights, with a 14 hr:10 hr day:night photoperiod and a 20°C/15 °C day/night temperature regime. We allowed 4 weeks of establishment for populations C1, C2, S1, and S2, and 5 weeks of establishment for populations N1 and N2 due to differences in germination time and growth rate between populations (Wooliver and Sheth, personal observation). Trays were sub-irrigated with a general nutrient solution (containing NPK + 13 micronutrients) three times per week (Mondays, Wednesdays, Fridays). Twice per week (Tuesdays, Thursdays), we emptied any excess water out of the trays to limit the growth of fungus, algae, and gnat larvae in the soil, added supplemental water to areas that had dried out, and rotated trays to minimize positional effects.

The 24 trays were then divided into four groups of six (with each group containing two trays from each of the three regions). Each group of six trays was then transferred into one of four reach-in growth chambers (Percival LT-105X, Percival Scientific, Inc., Perry, Iowa, USA) for the heatwave experiment. Two chambers were assigned the heatwave treatment and two chambers were assigned as controls, with the six trays positioned randomly within each chamber. This design ensured that each seed family within each population and cohort combination was replicated four times in each temperature regime, for a total of 1,728 plants in the full experiment. Given the early growth stage of the plants used in the experiment (∼4-5 weeks from planting), temperature regimes mimicked the natural climate from early in the growing season (∼May 15 to June 15), with the aim of simulating the most extreme heatwave that occurred between ancestral and descendant cohorts among all source populations. Examination of absolute maximum temperature and extreme daily temperature (95th percentile of maximum) data from the six focal populations from 1991-2023 (PRISM Climate Data; PRISM Group, 2014) guided our design of control and heatwave temperature regimes (Perkins and Alexander, 2013; Breshears et al., 2021; **Figure 1**). Duration of temperatures above the 95th percentile of maximum temperature in these populations ranged from 2-9 days, leading to the selection of a 5-day heatwave treatment, while the absolute maximum temperature from population S2 (37.7°C; June 4, 2016) guided the temperature regime for the heatwave treatment. Heatwave chambers were set to a 22°C/8°C day/night temperature regime for the first 3 days of the experiment to allow for plant acclimation before the 5-day heatwave treatment was applied, with a temperature regime of 38°C/14°C day/night and a 14 hr:10 hr day:night photoperiod (with a light intensity of ∼477 µmol/m²/s during the day period). Control chambers were set to a 22°C/8°C day/night temperature regime for the entire 8-day duration of the experiment. Temperature was ramped up and ramped down over the first 4 hours and last 4 hours of the day period, at a rate of 2.8°C/hr for the 22°C/8°C day/night temperature regime and 4.8°C/hr for the 38°C/14°C day/night temperature regime. Light intensity was ramped up and ramped down over the first 2 hours and last 2 hours of the day period at a rate of ∼20% of the daytime value per hr. The experiment was conducted between March 25 to April 11, 2024, with some staggering between chambers that matched the staggered planting dates of each group of six trays, which ensured consistent development stage across all trays within each growth chamber run. During the heatwave experiment, trays were sub-irrigated with reverse osmosis water rather than nutrient solution to eliminate confounding effects if higher temperatures increased plant water use.

### Focal traits

We measured six traits as important indicators of the ability of plants to respond to extreme heat while maintaining vital performance: *g*_sw_ (measured in mmol m^−2^ s^−1^), leaf temperature difference, Φ_PSII_, RGR in leaf number (hereafter referred to simply as RGR), SLA, and LDMC. On the final day of the heatwave (day 8), we used a handheld porometer/fluorometer (LI-600, LI-COR Biosciences, Lincoln, NE, USA) to measure *g*_sw_, leaf temperature difference, and Φ_PSII_, using the youngest fully-expanded leaf at the second or third node and excluding plants that were too small to fill the aperture of the porometer. *g*_sw_ increases with temperature due to the temperature dependence of diffusivity of water vapor through stomata (Turner, 1991; Urban et al., 2017; Griffani et al., 2024; Mills et al., 2024), which could confound our results when comparing *g*_sw_ between control and heatwave chamberss. Therefore, we also calculated the diffusion coefficient of water vapor (*D*_wv_) for each stomatal conductance measurement based on the air temperature in the growth chamber at the time of measurement (to the nearest minute) and assuming constant pressure as:

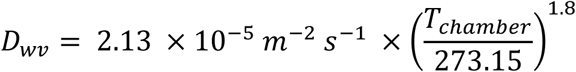

using 2.13 × 10^−5^ m^−2^ s^−1^ as the baseline diffusion coefficient for water vapor at 0°C and 1 atm of pressure (Nobel, 2020). For each control plant, we were then able to calculate the expected increase in *g*_sw_ based solely on a temperature increase from the air temperature in the control chamber to the heatwave chamber temperature of 38°C as:

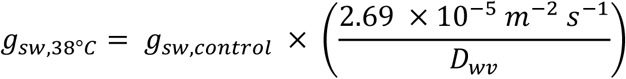

using 2.69 × 10^−5^ m^−2^ s^−1^ as the diffusion coefficient for water vapor at 38°C and 1 atm of pressure (Nobel, 2020), the diffusion coefficient for water vapor for each control plant calculated above, and the actual *g*_sw_ data for each control plant.

To measure RGR, we counted all true leaves > 1 mm in length on day 3, one day prior to initiation of the heatwave treatment (leaf_in_), and on day 9, the day after the heatwave treatment concluded (leaf_out_). To quantify the performance of seedlings, as the relative change in leaf number over the course of the heatwave treatment, we calculated RGR as: (leaf_out_ − leaf_in_)/(leaf_in_ × # of days). Although RGR is not a direct measure of reproductive output (i.e. fitness), size is positively correlated with fruit number in natural populations (Sheth and Angert, 2018) and has been used in past studies of *M. cardinalis* as a metric of relative performance (Wooliver et al., 2020). The day after the heatwave treatment concluded (day 9), we also harvested the youngest fully-expanded leaf at the second or third node from each plant. We measured leaf wet mass with a microbalance (MyWeigh iBalance 311, HBI International, Phoenix, AZ, USA) and leaf area with a bench-top leaf area meter (LI-3100C, LI-COR Biologicals, Lincoln, NE, USA). Leaves were then dried at 65°C for a minimum of one week and weighed with a microbalance (XSR104, Mettler-Toledo, Columbus, OH, USA) to determine dry mass. We calculated SLA as wet leaf area/dry leaf mass (measured in cm²/g). We calculated LDMC as dry leaf mass/wet leaf mass (measured in mg/g). In addition, we measured the length of the longest leaf prior to the initiation of the heatwave treatment as a covariate to account for differences in initial size at the start of the experiment. However, length of the longest leaf had no impact on how any independent variable affected any of the measured traits and so it was excluded from the final models.

### Statistical analyses

All statistical analyses were conducted in R Version 4.5.0 (2025) and R Studio Version 2024.12.1 (R Development Core Team, 2025). The six traits described above serve as our response variables of interest. SLA was log-transformed, while LDMC was square-root-transformed in order for analyses to meet assumptions of normality and homogeneity of variance.

To analyze physiological traits (*g*_sw_, leaf temperature difference, and Φ_PSII_), we constructed a model including fixed effects of cohort, region, heatwave treatment, and all two– and three-way interactions. Family was included as a random effect to account for non-independence of replicates with each family. We included time of day as a covariate because of its expected effects on these three physiological traits of interest and because of inconsistencies in the order in which individuals were selected for physiological trait measurement that could result in confounding of our independent variables of interest. We also included some of the possible two-way interactions between time of day and each fixed effect of interest, as well as three-way interactions between time of day, region, and heatwave treatment to further account for this potential for confounding. However, we avoid direct interpretations of the main effect of time of day or any interactions involving time of day and we do not anticipate any confounding of independent variables of interest based on time of day to affect our conclusions. A two-way interaction between region (leading edge versus range center versus trailing edge) and the heatwave treatment (heatwave versus control) would indicate variation in physiological, performance, and functional trait response to extreme heat across the range of *M. cardinalis* (objective 1). A two-way interaction between cohort (ancestors versus descendants) and the heatwave treatment would indicate an evolutionary response to the recent extreme heat event that affects how *M. cardinalis* responds to high temperatures (objective 2). A two-way interaction between cohort and region would indicate divergent evolutionary responses to the recent extreme heat event across the range of *M. cardinalis* (objectives 1 and 2). A three-way interaction between cohort, region, and the heatwave treatment would indicate divergent evolutionary responses across the range of *M. cardinalis* that affect how different populations across the range might respond to future extreme heat events.

To analyze RGR, SLA, and LDMC, we constructed a similar model to the one described above, but only including the fixed effects of heatwave treatment, cohort, region, and all two– and three-way interactions, along with family as a random effect, because time of day is not expected to affect these variables. We also included similar models, replacing region with population, in the Supplementary Material for completeness (**Table S1**). In general, the two source populations within each region responded in relatively similar ways to the heatwave treatment and between cohorts (**Figure S1**; **Figure S2**).

All general linear mixed effects models were conducted using the *glmmTMB* function in the *glmmTMB* package (version 1.1.5; (Brooks et al., 2017). All test statistics and *P*-values were calculated using the *Anova* function in the *car* package (version 3.1-2; (Fox and Weisberg, 2019) with Type II sums-of-squares. The model for leaf temperature difference resulted in significant autocorrelation with time of day, so we used the *gls* function in the *nlme* package (version 3.1-168; Pinheiro et al., 2021) and the *dispformula* argument in the *glmmTMB* function to build a more accurate model to account for this autocorrelation. For all significant interactions, estimated marginal means, custom contrasts, and effect sizes used to compare slopes between groups were generated using the *emmeans* and *contrast* functions in the *emmeans* package (version 1.10.4; (Lenth, 2025). This approach does often lead to substantial shrinking of effects sizes, particularly for the transformed variables; however, despite this, all trends reported below reflect statistically significant differences.

## Results

Region, heatwave treatment, and their two-way interaction affected *g*_sw_ (**Table 1**). Heatwave-treated plants exhibited greater *g*_sw_ than control plants (100%). When exposed to the heatwave treatment, trailing-edge plants experienced greater increases in *g*_sw_ than both range-center and leading-edge plants (151%, 86%, and 67% increases, respectively; **Table 1**, **S2**; **Figure 2A**). However, *g*_sw_ did not differ between cohorts and cohort did not exhibit any two-or three-way interactions with region and/or heatwave treatment to affect *g*_sw_ (**Table 1**).

**Figure 2.**
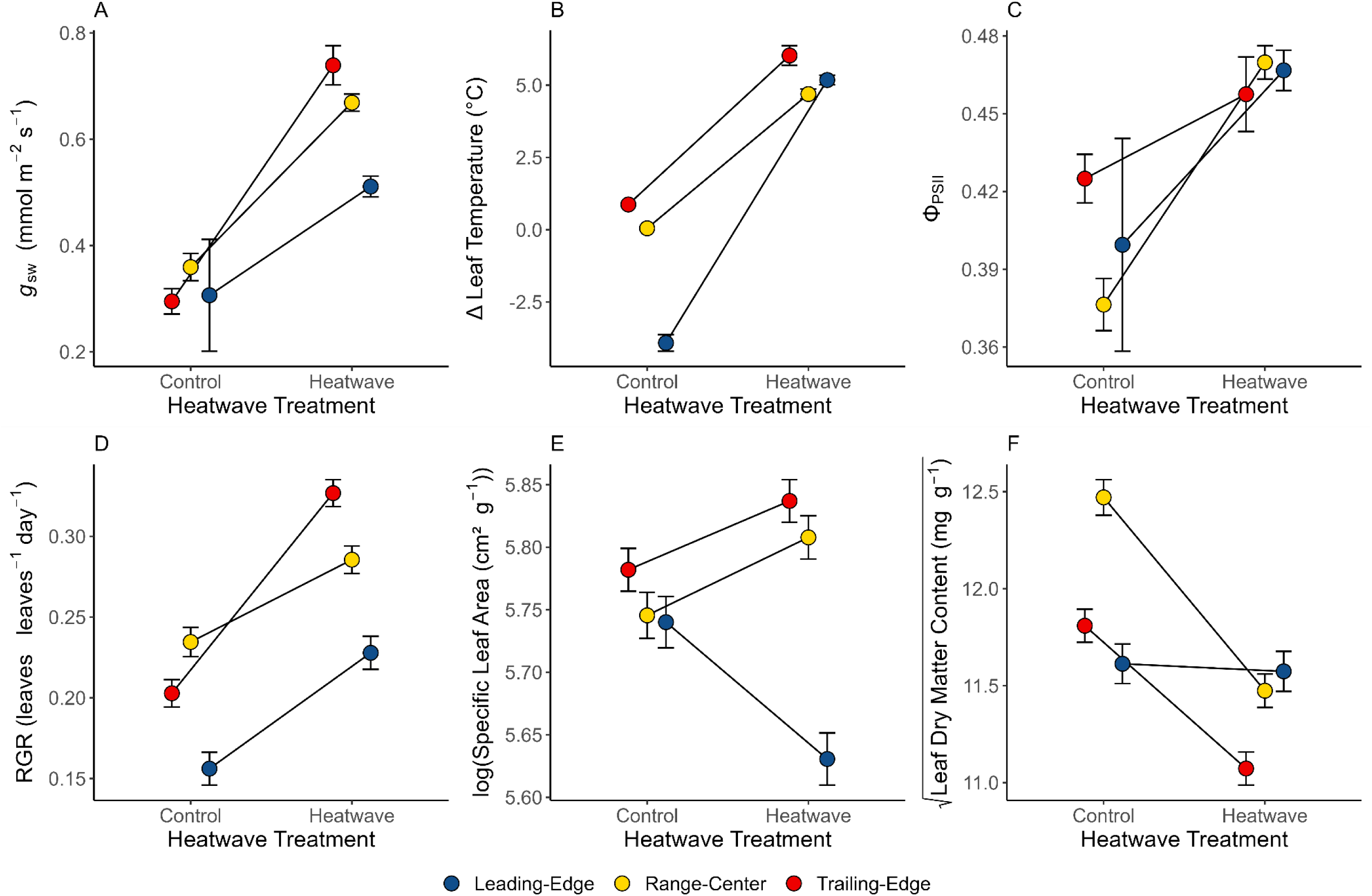
The effect of the heatwave treatment on physiological, performance, and functional traits of *M. cardinalis* plants from the three regions (leading edge, range center, and trailing edge) within its range. Measured traits include *g*_sw_ (A), leaf temperature difference (B), Φ_PSII_ (C), RGR (D), SLA (E), and LDMC (F). Error bars are ± one standard error around each estimated marginal mean.

**Table 1.** Results of mixed effects models assessing how cohort (2010 ancestors or 2017 descendants), region (leading edge, range center, or trailing edge), and heatwave treatment (treated or control) and their interactions affected *M. cardinalis* functional traits (*g*_sw_, Δ leaf temperature, Φ_PSII_, RGR, SLA, and LDMC). Post-hoc slope contrasts provide statistical tests of differences in slope among regions when comparing ancestors versus descendants or when comparing control plants versus heatwave treated plants, with the sign being in reference to the first half of the listed contrast. Significant effects and slope contrasts (*P* < 0.05) are listed in bold.

| | $g_{sw}$ | | | $\Delta$ Leaf Temperature | | | $\Phi_{PSII}$ | | |
| --- | --- | --- | --- | --- | --- | --- | --- | --- | --- |
| Predictor | $\chi^2$ | df | $P$ | $\chi^2$ | df | $P$ | $\chi^2$ | df | $P$ |
| Cohort (C) | 2.344 | 1 | 0.126 | 0.051 | 1 | 0.821 | 0.046 | 1 | 0.830 |
| Region (R) | **37.529** | **2** | **< 0.001** | **1832.65** | **2** | **< 0.001** | **58.183** | **2** | **< 0.001** |
| Heatwave (H) | **430.297** | **1** | **< 0.001** | **5701.89** | **1** | **< 0.001** | **447.035** | **1** | **< 0.001** |
| Time (T) | 1.371 | 1 | 0.242 | **302.324** | **1** | **< 0.001** | **29.583** | **1** | **< 0.001** |
| C × R | 0.306 | 2 | 0.858 | 1.991 | 2 | 0.370 | 0.886 | 2 | 0.642 |
| C × H | 3.011 | 1 | 0.083 | **5.705** | **1** | **0.017** | 0.322 | 1 | 0.571 |
| R × H | **16.704** | **2** | **< 0.001** | **198.774** | **2** | **< 0.001** | **51.360** | **2** | **< 0.001** |
| R × T | 2.114 | 2 | 0.348 | **94.469** | **2** | **< 0.001** | 4.031 | 2 | 0.133 |
| H × T | **9.322** | **1** | **0.002** | **166.083** | **1** | **< 0.001** | **30.150** | **1** | **< 0.001** |
| C × R × H | 2.346 | 2 | 0.309 | 0.812 | 2 | 0.666 | 0.785 | 2 | 0.675 |
| R × H × T | 1.751 | 2 | 0.417 | **9.213** | **2** | **< 0.001** | **50.191** | **2** | **< 0.001** |
| *Pairwise Slope Contrasts* |  |  |  |  |  |  |  |  |  |
| (Region × Heatwave) | Sign | | $P$ | Sign | | $P$ | Sign | | $P$ |
| Leading vs. Center | = |  | 0.346 | > |  | **< 0.001** | = |  | 0.521 |
| Leading vs. Trailing | < |  | **0.039** | > |  | **< 0.001** | = |  | 0.510 |
| Center vs. Trailing | < |  | **0.012** | = |  | 0.187 | > |  | **0.006** |

|  | RGR in Leaf Number |  |  | Specific Leaf Area |  |  | Leaf DMC |  |  |
| --- | --- | --- | --- | --- | --- | --- | --- | --- | --- |
| Predictor | $\chi^2$ | df | $P$ | $\chi^2$ | df | $P$ | $\chi^2$ | df | $P$ |
| Cohort (C) | 0.281 | 1 | 0.596 | 1.572 | 1 | 0.041 | 1.780 | 1 | 0.182 |
| Region (R) | **70.691** | **2** | **< 0.001** | **44.150** | **2** | **< 0.001** | **34.864** | **2** | **< 0.001** |
| Heatwave (H) | **130.600** | **1** | **< 0.001** | 0.933 | 1 | 0.334 | **79.103** | **1** | **< 0.001** |
| C × R | 3.639 | 2 | 0.162 | **19.789** | **2** | **< 0.001** | **12.489** | **2** | **0.002** |
| C × H | 0.002 | 1 | 0.966 | 1.137 | 1 | 0.286 | 0.474 | 1 | 0.491 |
| R × H | **18.996** | **2** | **< 0.001** | **24.315** | **2** | **< 0.001** | **27.236** | **2** | **< 0.001** |
| C × R × H | 0.016 | 2 | 0.992 | 0.077 | 2 | 0.962 | 0.506 | 2 | 0.776 |
| *Pairwise Slope Contrasts* |  |  |  |  |  |  |  |  |  |
| (Cohort × Region) | Sign | | $P$ | Sign | | $P$ | Sign | | $P$ |
| Leading vs. Center | — |  | — | = |  | 0.602 | = |  | 0.934 |
| Leading vs. Trailing | — |  | — | > |  | **0.001** | < |  | **0.003** |
| Center vs. Trailing | — |  | — | > |  | **< 0.001** | < |  | **0.002** |
| *Pairwise Slope Contrasts* |  |  |  |  |  |  |  |  |  |
| (Region × Heatwave) | Sign | | $P$ | Sign | | $P$ | Sign | | $P$ |
| Leading vs. Center | = |  | 0.277 | < |  | **< 0.001** | > |  | **< 0.001** |
| Leading vs. Trailing | < |  | **0.005** | < |  | **< 0.001** | > |  | **< 0.001** |
| Center vs. Trailing | < |  | **< 0.001** | = |  | 0.834 | = |  | 0.123 |

Region, heatwave treatment, their two-way interaction, and a two-way interaction between cohort and heatwave affected leaf temperature difference (air temperature − leaf temperature; **Table 1**). Heatwave-treated plants exhibited lower leaf temperatures than air temperatures (5.3°C lower, on average), while control plants exhibited higher leaf temperatures than air temperatures (0.7°C higher, on average). Overall, leaf temperature differences were strongly affected by the earlier measurement time of day for many leading-edge control plants, so it is difficult to make meaningful conclusions from slope contrasts based on the cohort × heatwave and region × heatwave interactions, so we limit our interpretation to only heatwave-treated plants. When exposed to the heatwave treatment, ancestors and descendants exhibited similar leaf temperature differences (5.1°C, and 5.4°C, respectively), while trailing-edge plants exhibited larger average leaf temperature differences than leading-edge and range-center plants (raw means of 6.2°C, 4.6°C, and 4.8°C, respectively; **Table 1**, **S2**; **Figure 2B**). Leaf temperature differences were similar between cohorts and cohort did not exhibit a two-way interaction with region or a three-way interaction with region and heatwave treatment to affect leaf temperature difference (**Table 1**).

Region, heatwave treatment, and their two-way interaction affected Φ_PSII_ (**Table 1**). Heatwave-treated plants exhibited greater Φ_PSII_ than control plants (8%). When exposed to the heatwave treatment, range-center plants experienced greater increases in Φ_PSII_ than both leading-edge and trailing-edge plants (13%, 8%, and 4% increases, respectively; **Table 1**, **S2**; **Figure 2C**). Φ_PSII_ did not differ between cohorts and cohort did not exhibit any two-or three-way interactions with region and/or heatwave treatment to affect Φ_PSII_ (**Table 1**).

Region, heatwave treatment, and their interaction affected RGR (**Table 1**). Heatwave-treated plants exhibited greater RGR than control plants (41%). When exposed to the heatwave treatment, trailing-edge plants experienced larger increases in RGR than leading-edge and range-center plants (61%, 46%, and 22% increases, respectively; **Table 1**, **S2**; **Figure 2D**). RGR did not differ between cohorts and cohort did not exhibit any two-or three-way interactions with region and/or heatwave treatment to affect RGR (**Table 1**).

Region and two-way interactions between region and either cohort or heatwave treatment affected SLA (**Table 1**). When comparing descendants to ancestors, leading edge and range-center plants increased in SLA (both 1%), while trailing-edge plants decreased in SLA (1%; **Table 1**, **S2**; **Figure 3A**). When exposed to the heatwave treatment, range-center and trailing-edge plants increased in SLA (both 1%), while leading edge plants decreased (2%; **Table 1**, **S2**; **Figure 2E**). SLA did not differ between cohorts or by heatwave treatment, and heatwave treatment did not interact with cohort or both cohort and region to affect SLA (**Table 1**).

**Figure 3.**
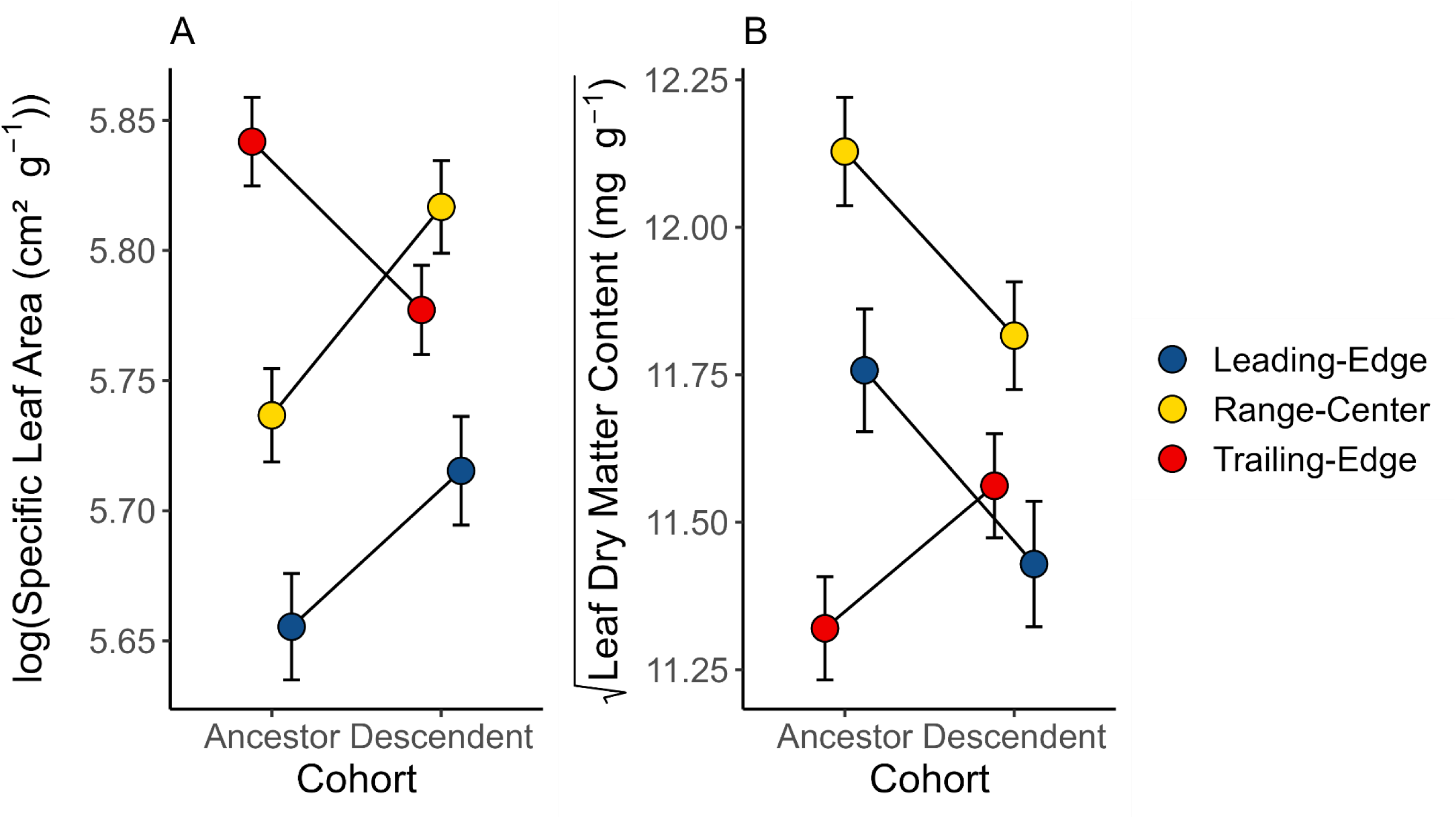
The effect of cohort (ancestors versus descendants) on functional traits of *M. cardinalis* plants from the three regions (leading edge, range center, and trailing edge) within its range. Measured traits include SLA. (A) and LDMC (B). Error bars are ± one standard error around each estimated marginal mean.

Region, heatwave treatment, and two-way interactions between region and either cohort or heatwave treatment affected LDMC (**Table 1**). Heatwave-treated plants exhibited lower LDMC than control plants (5%). When comparing descendants to ancestors, leading-edge and range-center plants decreased in LDMC (3% and 2%, respectively), while trailing-edge plants increased (3%; **Table 1**, **S2**; **Figure 3B**). When exposed to the heatwave treatment, range-center and trailing-edge plants exhibited decreases in LDMC (8%, and 6%, respectively), while leading-edge plants were not affected (**Table 1**, **S2**; **Figure 2F**). LDMC did not differ between cohorts, and heatwave treatment did not interact with cohort or both cohort and region to affect LDMC (**Table 1**).

## Discussion

We investigated responses to a repeated heatwave event in populations from across the range of *M. cardinalis* by testing for changes in plant physiological, performance, and functional traits of resurrected ancestor (pre-heatwave) and descendant (post-heatwave) populations exposed to a heatwave treatment. Heatwave-treated plants demonstrated greater plant physiological performance (increased *g*_sw_ and Φ_PSII_), along with faster development of cheaper leaves (increased RGR and decreased LDMC), compared to control plants, which was contrary to our predictions and indicative of a lack of heat stress. Responses to the heatwave treatment also varied among regions, with populations from the trailing edge of the range of *M. cardinalis* exhibiting the greatest positive response for *g*_sw_ and RGR, but not for Φ_PSII_, while SLA and LDMC diverged among regions in response to extreme heat. Ancestors and descendants did not differ in any measured traits, overall, potentially indicating a lack of evolutionary response to the recent heatwave in western North America. When evaluated between regions, however, there was some evidence of divergent evolutionary responses to extreme climate events. Namely, when exposed to the heatwave treatment, trailing-edge and range-center populations developed larger, but cheaper leaves, while leading-edge populations developed smaller leaves that were more expensive per unit area, relative to the control treatment. Ancestor and descendant cohorts also did not exhibit divergent responses to the heatwave treatment, indicating a lack of evidence of adaptive evolution that may increase the ability of populations to withstand heatwaves. Here we discuss our results with respect to both plant-level trait responses and population-level evolutionary responses to extreme heat events.

### Plant trait responses to heatwave treatment

We observed an unexpected, strong, and positive response to the heatwave treatment in terms of increased *g*_sw_ and Φ_PSII_, along with increased growth rate of cheaper leaves. In general, heat stress causes stomatal closure and inhibits photosynthetic enzymes, so it is unlikely that the heatwave treatment was severe enough to elicit a stress response (Hasanuzzaman et al., 2013; Feller and Vaseva, 2014; Teskey et al., 2015; dos Santos et al., 2022). Instead, in the absence of stress-induced stomatal closure, *g*_sw_ will increase with temperature both due to the physical effects of temperature-dependent diffusivity, as well as plant level responses to a rise in temperature, such as increased stomatal density or stomatal opening (Turner, 1991; Urban et al., 2017; Griffani et al., 2024; Mills et al., 2024). Overall, we calculated the physical effects of temperature-dependent diffusivity to be responsible for ∼13% of the increase in *g*_sw_ for heatwave-treated plants (**Figure S3**), demonstrating that plant physiological responses to the temperature increase are predominantly responsible for the increased performance for heatwave-treated plants in terms of gas exchange (Hasanuzzaman et al., 2013; dos Santos et al., 2022). Increased *g*_sw_ also provides a cooling effect to leaves that could prevent heat stress from occurring (Farquhar and Sharkey, 1982; Teskey et al., 2015; Griffani et al., 2024). In our experiment, control growth chambers were set to a daytime temperature of 22°C and the plants therein had an average leaf temperature of 21.4°C, while heatwave chambers were set to a daytime temperature of 38°C and the plants therein had an average leaf temperature of 31.6°C, demonstrating a substantial cooling effect. It is therefore possible that leaf cooling through stomatal conductance (i.e., latent heat loss through transpiration) allowed heatwave-treated plants to avoid negative physiological effects of high temperatures, while still allowing photosynthetic machinery to operate at a faster rate compared to control plants that were maintained at a temperature that was below optimal in controlled conditions (ambient CO_2_ and sufficient light; Farquhar and Sharkey, 1982; Teskey et al., 2015; Griffani et al., 2024).

Within the general trend of increased performance for heatwave-treated plants, individuals from the trailing edge of the range of *M. cardinalis* often experienced the most substantial performance increase relative to the control treatment. The relatively high increase in *g*_sw_ and RGR in trailing-edge plants is consistent with the natural environment in which these source populations are found. In general, trailing-edge populations (in our study, population S2, specifically; **Table S1**) have experienced historic and recent climate conditions that are warmer than those of range-center and leading-edge populations, which led to the prediction that these populations would exhibit greater resistance (i.e., a less negative response) to the heatwave treatment. Although the overall positive response to the heatwave treatment across the whole experiment was not in line with our predictions, observing the greatest positive response for trailing-edge plants is not unexpected (Angert et al., 2011; King et al., 2019; Chiono and Paul, 2023). Conversely, leading-edge populations have been shown to exhibit greater plasticity of thermally-regulated gene expression, which could reduce their vulnerability to future climate fluctuations (Preston et al., 2022). In addition, trailing-edge populations of *M. cardinalis* often exhibit a more annualized life history strategy, with rapid, early growth and overall greater vegetative investment (i.e., variation that was captured by our study of responses to a short-term heatwave by plants at an early development stage; (Nelson et al., 2021). By contrast, heatwaves in southern California often occur late in the growing season, possibly having a greater impact on reproductive output than on early season vegetative growth. A future study involving experimental plants grown for an entire reproductive cycle (including a measure of reproductive fitness) and subjected to a heatwave of greater intensity or duration would provide more definitive insight into different trends for leading-edge, range-center, and trailing-edge population responses to heatwave treatments.

### Environment-specific evolutionary responses to a recent heatwave event

Across all measured traits, we observed a lack of overall differences between ancestors and descendants. This is not unexpected based on measures of heritability in some of the same traits we examined from a field experiment using the same six *M. cardinalis* source populations as the present study (Sheth et al., in review), indicating limited evolutionary potential within these populations. However, it is important to highlight that an overall effect of cohort on SLA and LDMC could have been masked by descendants from each region responding in different directions compared to ancestors, with trailing-edge descendants trending towards smaller, more expensive leaves and leading-edge and range-center descendants trending towards larger, cheaper leaves. In general, smaller and thicker leaves tend to be selected for in hotter environments (i.e. the trailing edge), while populations that are less adapted to high temperatures may opt for more conservative resource investment strategies, which is in line with our results (Knight and Ackerly, 2003; Reich, 2014; Leigh et al., 2017; Liu et al., 2023). However, the focal *M. cardinalis* populations all experienced not only extreme heat, but also extreme drought from 2010 to 2017, which can also play a substantial role in selection on plant traits, resource investment, and thermal tolerance (Feller and Vaseva, 2014; Teskey et al., 2015; Anstett et al., 2021; Cook et al., 2021; dos Santos et al., 2022). All experimental plants in this study were kept well-watered, so it is difficult to identify the relative contributions of extreme temperature and soil moisture conditions on plant traits in order to make broad conclusions regarding the evolutionary response to extreme climate events in this system.

The heatwave treatment was also unsuccessful in eliciting a heat stress response in our experimental plants, which could have masked our ability to detect an evolutionary response between ancestor and descendant cohorts (i.e., both cohorts experienced substantially higher performance in the heatwave treatment). In fact, thermal performance curves for the same source populations as the present study predict greater performance (in terms of RGR) at 38°C than at 22°C for all populations (Wooliver et al., 2020). This result is consistent with classic thermal performance curve hypotheses, which posit that organisms are optimally placed at “suboptimal” temperatures on a thermal performance curve, due to its left-skewed asymmetry, which results in greater risk of large declines in fitness at relatively modest temperature increases compared to lower risk at similarly modest temperature decreases (Martin and Huey, 2008). Additionally, past thermal performance curves for our *M. cardinalis* source populations also indicate that ancestral cohorts (particularly for range-center and leading-edge populations) had thermal optima that were warmer than their historic maximum July temperatures (Wooliver et al., 2020). This could indicate that *M. cardinalis* populations have already adapted to high yearly average and extreme temperatures prior to the ancestral 2010 cohort (**Table S1**). Additionally, recent heatwaves may have not exposed *M. cardinalis* populations to temperatures that extend above the upper range of their thermal performance breadth, reducing the likelihood of a strong evolutionary response to those climate extremes. However, as temperatures continue to rise and become closer to T_50_, the thermal safety margin of evolutionarily static *M. cardinalis* populations will become smaller, indicating the potential for future vulnerability to high temperatures (i.e., a lack of evolutionary rescue in response to climate change; (Wooliver et al., 2020; Kitudom et al., 2022). Alternatively, if substantial adaptation to warming (or extreme heat) occurs in response to both recent and future heatwaves, thermal safety margins of plant populations could rise, increasing the likelihood of evolutionary rescue (Ahrens et al., 2021).

Thermal optima in controlled conditions may not be representative of the climate optima for these populations in their natural environments, which are likely to be less predictable due to natural environmental variability and the involvement of numerous additional environmental variables. For example, water availability for riparian species like *M. cardinalis* is likely to be more transient throughout the growing season, while plants in this experiment remained well-watered throughout the entire heatwave treatment, removing the potential for negative effects of stomatal opening on water loss under drought conditions (Cook et al., 2021; Posch et al., 2024). Additionally, the heatwave treatment may have been mistimed with what is likely to be experienced by plants at the particular developmental stage of those used in the experiment. A future experiment simultaneously manipulating extreme heat and drought and/or testing the effects of various heatwave treatments could help elucidate how plant populations will respond to realistic extreme conditions that are likely to increase in intensity, frequency, duration, and geographic extent over time (Meehl and Tebaldi, 2004; Seneviratne et al., 2012; Coumou and Robinson, 2013; Coumou et al., 2013; Perkins-Kirkpatrick and Gibson, 2017; Guerreiro et al., 2018; IPCC, 2022).

### Conclusions

Extreme climate events can play an important role in imposing selection in natural populations and could increasingly threaten the maintenance of biodiversity if populations are unable to cope with their frequency, duration, and intensity in the future. Overall, we found limited differences between ancestral and descendant cohorts of *M. cardinalis* harvested before and after a repeated heat and drought event in western North America, restricting our ability to make predictions about how *M. cardinalis* populations will respond to future extreme heat events. However, we did find divergent trends in SLA and LDMC between cohorts for populations from different areas of the range of *M. cardinalis*, indicating that evolutionary responses to extreme climate are likely to be environment-specific. We also found that across numerous traits, plant performance increased for heatwave treated plants and often increased most substantially in plants from the warmest local environments at the trailing edge of the range. This result indicates that either recent heatwaves have not been extreme enough to impose strong selection on *M. cardinalis* populations, potentially mediated by historical adaptive evolution to extreme heat and/or heat tolerance strategies, such as leaf cooling through stomatal conductance, leaf angle, or various molecular mechanisms (Hasanuzzaman et al., 2013; dos Santos et al., 2022). Many *M. cardinalis* populations therefore likely reside in natural environments that are below their optimum average temperature in order to avoid modest temperature increases posing a substantial threat to survival and fitness. However, the expanding intensity, frequency, and duration of heatwaves, compounded with associated effects of drought, could result in sustained and/or repeated high temperatures that pose an increasing threat to performance and fitness if populations are unable to undergo evolutionary rescue, highlighting the importance of continued study of the effects of extreme climate events on natural populations.

## Supporting information

Albano_et_al_2025_SupplementaryMaterial

## Acknowledgements

We thank the North Carolina State University Phytotron staff, especially Deepti Pradhan, for their assistance with setting up, maintaining, and conducting the experiment. We also thank Dachuan Wang for advice on plant physiological measurements taken by the porometer. This work was funded by the National Science Foundation grant DEB-2131815 to SNS (including a Research Opportunity Award Supplement to support RAB) and DEB-2131817 to CDM, and Research Capacity Fund (HATCH), project award no. 7002993, from the U.S. Department of Agriculture’s National Institute of Food and Agriculture to SNS.

## Data Availability Statement

The dataset generated and analyzed during this study, along with the associated code for statistical analysis, is published in Dryad Digital Repository (https://doi.org/10.5061/dryad.905qfttx8).

## Notes

https://doi.org/10.5061/dryad.905qfttx8

### Competing Interest Statement

The authors have declared no competing interest.

