## Supplementary material for "Range-wide responses to an extreme heat event in *Mimulus cardinalis*": Albano_et_al_2025_SupplementaryMaterial

**Table S1** Location and climate data for the six focal *M. cardinalis* populations used in this study. Location data includes latitude, longitude, and elevation for each population. Climate data includes mean temperature and max temperature (both limited to between May 15 and June 15 of each year) for a historical time period (1990-2010) and the recent climate extreme that is the focus of this study (2011-2017). All climate data was extracted from PRISM (PRISM Climate Data; (PRISM Group, 2014).

| | | | | Historical (1990-2010) | | Recent Climate Extreme (2011-2017) | |
|---|---|---|---|---|---|---|---|
| Population | Latitude (°N) | Latitude (°W) | Elevation (m) | Mean Temperature (05/15-06/15; °C) | Max Temperature (05/15-06/15; °C) | Mean Temperature (05/15-06/15; °C) | Max Temperature (05/15-06/15; °C) |
| N1 | 43.37876 | −122.95207 | 295 | 14.5 | 39.7 | 14.6 | 37.6 |
| N2 | 42.53529 | −123.73016 | 914 | 11.7 | 33.7 | 11.9 | 33.6 |
| C1 | 37.70377 | −119.75363 | 1316 | 14.9 | 34.1 | 15.0 | 35.0 |
| C2 | 37.54576 | −119.64152 | 1228 | 14.7 | 35.3 | 14.4 | 34.6 |
| S1 | 32.92788 | −116.56019 | 1252 | 14.6 | 33.7 | 14.7 | 31.6 |
| S2 | 32.60831 | −116.70098 | 252 | 18.3 | 39.7 | 18.2 | 37.7 |

**Table S2** Results of mixed effects models assessing how cohort (2010 or 2017), population (N1, N2, C1, C2, S1, or S2), and heatwave treatment (treated or control) and their interactions affected *M. cardinalis* physiological, performance, and functional traits ($g_{sw}$, Δ leaf temperature, $\Phi_{PSII}$, RGR, SLA, and LDMC). Three-way interactions were removed from the model explaining $\Phi_{PSII}$ due to convergence issues. Significant effects ($P$ < 0.05) are listed in bold.

| | $g_{sw}$ | | | Δ Leaf Temperature | | | $\Phi_{PSII}$ | | |
|---|---|---|---|---|---|---|---|---|---|
| Predictor | $\chi^2$ | df | *P* | $\chi^2$ | df | *P* | $\chi^2$ | df | *P* |
| Cohort (C) | 2.022 | 1 | 0.155 | 3.142 | 1 | 0.076 | 0.174 | 1 | 0.676 |
| Population (P) | **51.627** | **5** | **< 0.001** | **53.571** | **5** | **< 0.001** | **82.884** | **5** | **< 0.001** |
| Heatwave (H) | **441.739** | **1** | **< 0.001** | **3164.45** | **1** | **< 0.001** | **441.644** | **1** | **< 0.001** |
| Time (T) | 1.402 | 1 | 0.236 | **50.017** | **1** | **< 0.001** | **30.538** | **1** | **< 0.001** |
| C × P | 4.209 | 5 | 0.520 | 2.721 | 5 | 0.743 | 6.081 | 5 | 0.299 |
| C × H | 3.139 | 1 | 0.076 | 1.794 | 1 | 0.181 | 0.697 | 1 | 0.404 |
| P × H | **20.701** | **5** | **< 0.001** | **137.975** | **5** | **< 0.001** | **61.049** | **5** | **0.010** |
| P × T | 2.368 | 5 | 0.796 | 6.349 | 5 | 0.274 | 4.352 | 5 | 0.500 |
| H × T | **9.456** | **1** | **0.002** | **135.031** | **1** | **< 0.001** | **32.658** | **1** | **< 0.001** |
| C × P × H | 5.174 | 5 | 0.395 | 1.750 | 5 | 0.883 | — | — | — |
| P × H × T | 3.490 | 5 | 0.625 | **74.687** | **5** | **< 0.001** | — | — | — |

| | RGR in Leaf Number | | | Specific Leaf Area | | | Leaf DMC | | |
|---|---|---|---|---|---|---|---|---|---|
| Predictor | $\chi^2$ | df | *P* | $\chi^2$ | df | *P* | $\chi^2$ | df | *P* |
| Cohort (C) | 0.323 | 1 | 0.570 | 1.783 | 1 | 0.182 | 2.220 | 1 | 0.136 |
| Population (P) | **71.952** | **5** | **< 0.001** | **56.398** | **5** | **< 0.001** | **77.434** | **5** | **< 0.001** |
| Heatwave (H) | **130.951** | **1** | **< 0.001** | 0.821 | 1 | 0.365 | **78.817** | **1** | **< 0.001** |
| C × P | 3.677 | 5 | 0.597 | **19.864** | **5** | **0.001** | **13.727** | **5** | **0.017** |
| C × H | 0.004 | 1 | 0.953 | 1.139 | 1 | 0.286 | 0.633 | 1 | 0.426 |
| P × H | **19.612** | **5** | **< 0.001** | **28.851** | **5** | **< 0.001** | **30.861** | **5** | **< 0.001** |
| C × P × H | 1.235 | 5 | 0.942 | 0.493 | 5 | 0.992 | 0.812 | 5 | 0.976 |

**Table S3** Post-hoc slope contrasts and standard errors, used to provide the statistical tests of differences in slope (found in **Table 1**) between regions (leading edge, range center, or trailing edge) when comparing ancestors versus descendants (for significant Cohort × Region interactions) or comparing control plants versus heatwave treated plants (for significant Region × Heatwave interactions). Significant slopes ($P < 0.05$; **Table 1**) are listed in bold.

| | $g_{sw}$ | | Δ Leaf Temperature | | $\Phi_{PSII}$ | |
|---|---|---|---|---|---|---|
| *Pairwise Slope Contrasts (Region × Heatwave)* | Slope | SE | Slope | SE | Slope | SE |
| Leading vs. Center | − 0.105 | 0.111 | **4.270** | **0.712** | − 0.025 | 0.039 |
| Leading vs. Trailing | **− 0.240** | **0.116** | **5.700** | **0.741** | 0.027 | 0.041 |
| Center vs. Trailing | **− 0.135** | **0.053** | **1.430** | **0.341** | **0.052** | **0.019** |

| | RGR in Leaf Number | | Specific Leaf Area | | Leaf DMC | |
|---|---|---|---|---|---|---|
| *Pairwise Slope Contrasts (Cohort × Region)* | Slope | SE | Slope | SE | Slope | SE |
| Leading vs. Center | — | — | − 0.020 | 0.039 | − 0.016 | 0.197 |
| Leading vs. Trailing | — | — | **0.125** | **0.038** | **− 0.570** | **0.194** |
| Center vs. Trailing | — | — | **0.145** | **0.035** | **− 0.554** | **0.180** |
| *Pairwise Slope Contrasts (Region × Heatwave)* | Slope | SE | Slope | SE | Slope | SE |
| Leading vs. Center | 0.021 | 0.019 | **− 0.172** | **0.039** | **0.958** | **0.187** |
| Leading vs. Trailing | **− 0.052** | **0.019** | **− 0.164** | **0.035** | **0.697** | **0.183** |
| Center vs. Trailing | **− 0.073** | **0.017** | 0.007 | 0.038 | − 0.261 | 0.169 |

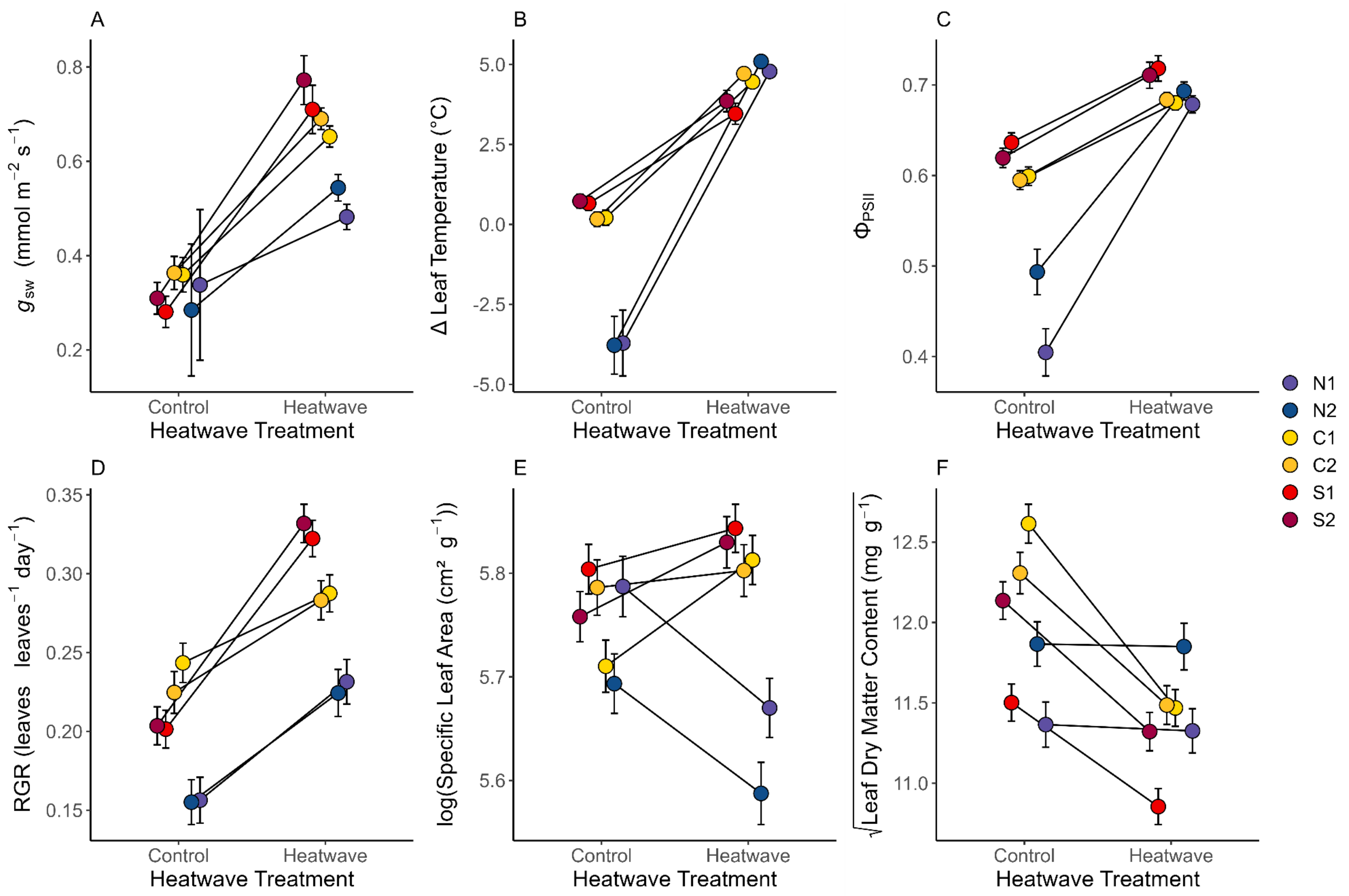
A
B
C
D
E
F
$g_{sw}$ (mmol m$^{-2}$ s$^{-1}$)
Δ Leaf Temperature (°C)
$\Phi_{PSII}$
RGR (leaves leaves$^{-1}$ day$^{-1}$)
log(Specific Leaf Area (cm² g$^{-1}$))
$\sqrt{\text{Leaf Dry Matter Content (mg g}^{-1})}$
0.2
0.4
0.6
0.8
-5.0
-2.5
0.0
2.5
5.0
0.4
0.5
0.6
0.7
0.15
0.20
0.25
0.30
0.35
5.6
5.7
5.8
11.0
11.5
12.0
12.5
Control
Heatwave
Heatwave Treatment
N1
N2
C1
C2
S1
S2

**Figure S1** The effect of the heatwave treatment on the physiological, performance, and functional traits of *M. cardinalis* plants from the two leading-edge (N1 and N2), two range-center (C1 and C2), and two trailing-edge (S1 and S2) source populations. Measured traits include $g_{sw}$ (A), leaf temperature difference (B), $\Phi_{PSII}$ (C), RGR (D), SLA (E), and LDMC (F). Error bars are ± one standard error around each estimated marginal mean.

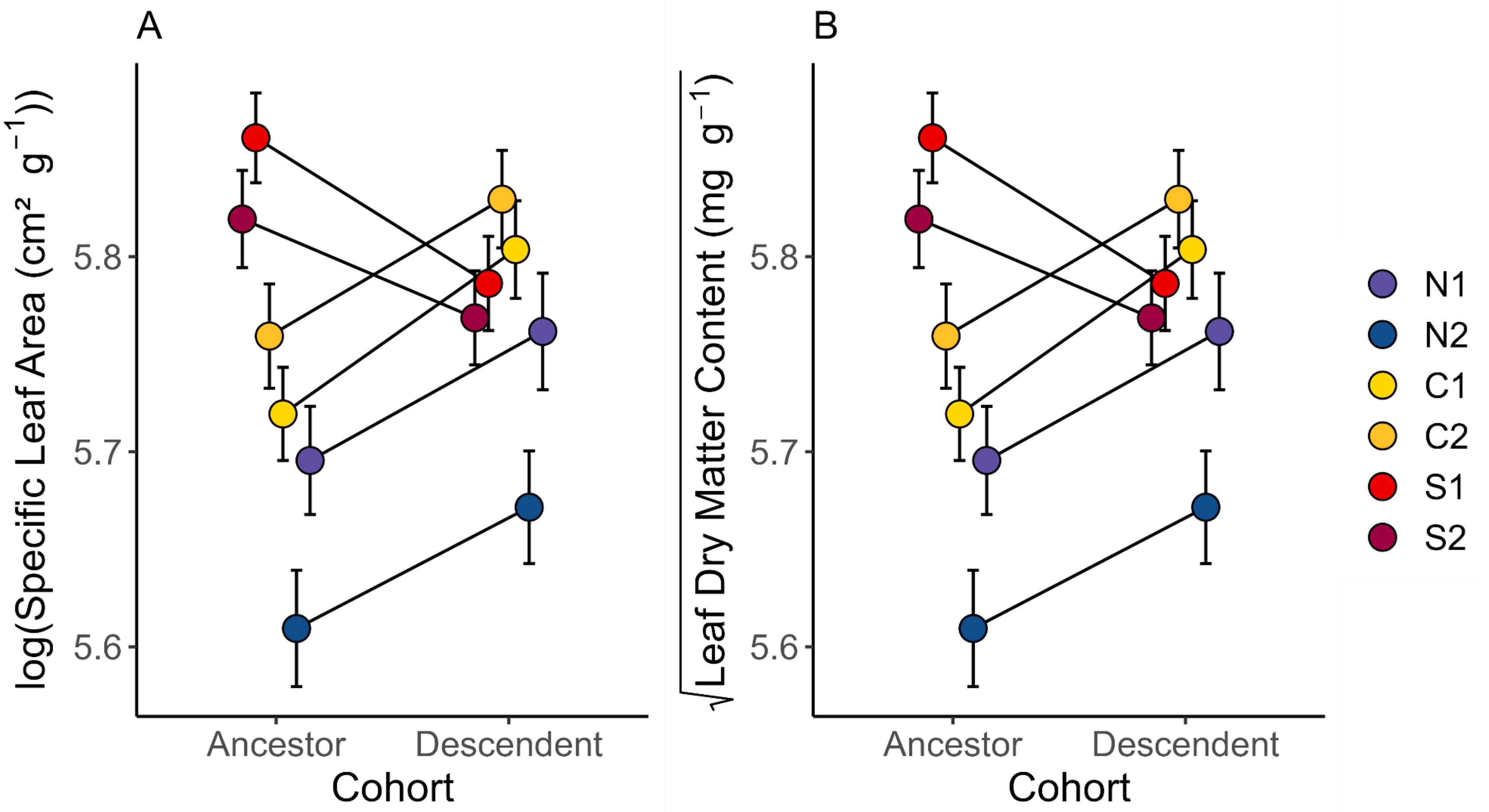


**Figure S2** The effect of cohort (ancestors versus descendants) on functional traits of *M. cardinalis* plants from the two leading-edge (N1 and N2), two range-center (C1 and C2), and two trailing-edge (S1 and S2) source populations. Measured traits include SLA (A) and LDMC (B). Error bars are ± one standard error around each estimated marginal mean.

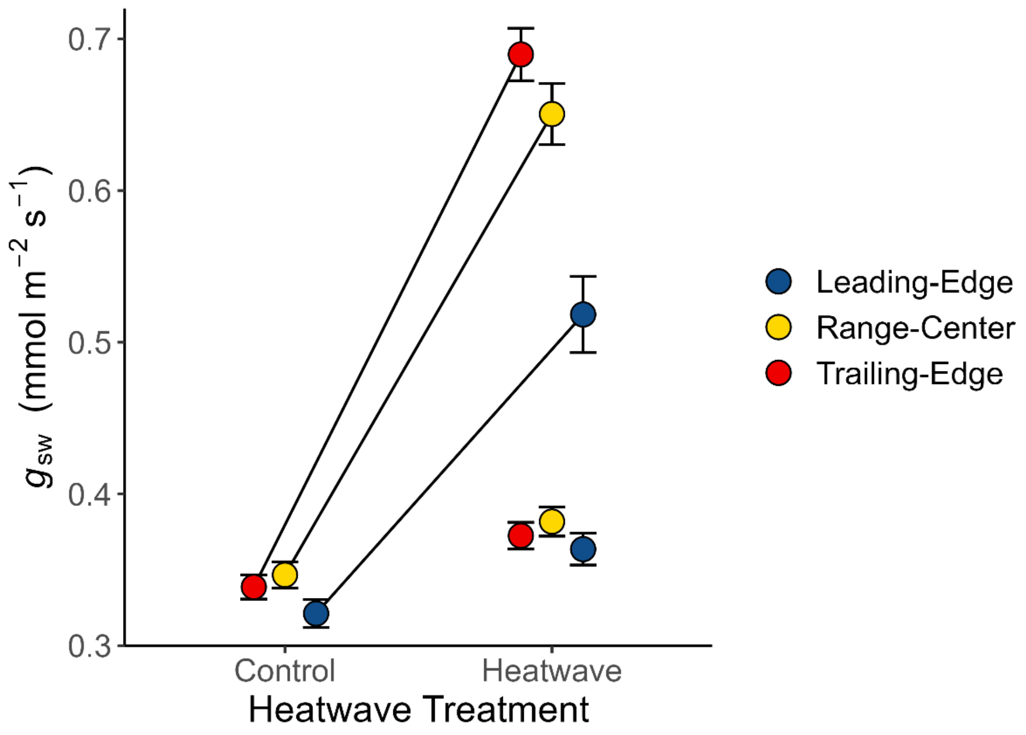


**Figure S3** The effect of the heatwave treatment on $g_{sw}$ of *M. cardinalis* plants from the three regions (leading edge, range center, and trailing edge) within its range. The additional set of points shows how control plants would be predicted to increase in $g_{sw}$ when exposed to the heatwave treatment, based solely on the expected temperature-dependent physical increase in diffusivity of water vapor through stomata, aside from any plant level response to extreme heat. Error bars are ± one standard error around each raw mean value.
